# New environment, new invaders - repeated horizontal transfer of LINEs to sea snakes

**DOI:** 10.1101/2020.02.27.968685

**Authors:** James D. Galbraith, Alastair J. Ludington, Alexander Suh, Kate L. Sanders, David L. Adelson

**Affiliations:** School of Biological Sciences, University of Adelaide, Adelaide, SA 5005, Australia; Department of Ecology and Genetics - Evolutionary Biology, Evolutionary Biology Centre, Uppsala University, SE-752 36 Uppsala, Sweden; Department of Organismal Biology - Systematic Biology, Evolutionary Biology Centre, Uppsala University, SE-752 36 Uppsala, Sweden; School of Biological Sciences, University of East Anglia, Norwich Research Park, NR4 7TU, Norwich, United Kingdom

**Keywords:** horizontal transfer, transposable element, Serpentes

## Abstract

While numerous studies have found horizontal transposon transfer (HTT) to be widespread across metazoans, few have focused on HTT in marine ecosystems. To investigate potential recent HTTs into marine species we searched for novel repetitive elements in sea snakes, a group of elapids which transitioned to a marine habitat at most 18 Mya. Our analysis uncovered repeated HTTs into sea snakes following their marine transition. Such major shifts in habitat should require significant genomic changes.The seven subfamilies of horizontally transferred LINE retrotransposons we identified in the olive sea snake (*Aipysurus laevis*) are transcribed, and hence are likely still active and expanding across the genome. A search of 600 metazoan genomes found all seven were absent from other amniotes, including terrestrial elapids, with the most similar transposons present in fish and marine invertebrates. The one exception was a similar transposon found in sea kraits, a lineage of amphibious elapids which independently transitioned to a marine environment following their divergence from terrestrial species 25 Mya. Our finding of repeated horizontal transfer events into separate lineages of marine snakes greatly expands past findings of frequent horizontal transfer in the marine environment, suggesting it is ideal for the transfer of transposons.Transposons are drivers of evolution as sources of genomic sequence and hence genomic novelty. This provides evidence of the environment influencing evolution of metazoans not only through specific selection pressures, but also by contributing novel genomic material.

## Introduction

Transposons are a major component of metazoan genomes, making up 8 to 28% of the typical amniote genome (Green et al. 2014; Zhang et al. 2014; Platt et al. 2018; Zeng et al. 2018). Transposons are split into two classes: Class I containing LINEs (long interspersed elements) and LTR (long terminal repeat) retrotransposons; and Class II containing DNA transposons (Wicker et al. 2007). Whilst transposons are normally vertically transmitted (parent to offspring) there have been many instances of horizontal transfer of transposons (HTT) observed between distantly related species. While HTT of DNA transposons and LTR retrotransposons appears to be more common, many examples of HTT of non-LTR retrotransposons (LINEs) have been described (Peccoud et al. 2018). These include transfers of RTE-BovBs between ticks and distant vertebrate lineages (Ivancevic et al. 2018), of AviRTEs between birds and parasitic nematodes (Suh et al. 2016), and of Rex1 elements between teleost fish (Volff et al. 2000; Zhang et al. 2020). As transposons proliferate throughout a genome they can contribute novel coding sequences, alter gene regulatory networks, modify coding regions and lead to gene copy number variation (Rebollo et al. 2012; Chuong et al. 2017; Cerbin and Jiang 2018; Schrader and Schmitz 2019). Within a lifetime most insertions will be neutral and some may be deleterious; however, on an evolutionary time scale, some TE insertions constitute a key source of genomic innovation as organisms adapt to new and changing environments (Casacuberta and González 2013; Salces-Ortiz et al. 2020).

Hydrophiinae (Elapidae) is a prolific radiation of more than 100 terrestrial snakes plus ∼70 aquatic species. The aquatic species form two separate lineages which independently transitioned to a marine habitat: the fully marine sea snakes and the amphibious sea kraits (*Laticauda*) (Lee et al. 2016). Sea snakes are phylogenetically nested inside the terrestrial hydrophiine radiation and appeared ∼6-18 Mya, while sea kraits form the sister lineage to all other Hydrophiinae and diverged 25 Mya (Lee et al. 2016; Sanders et al. 2008). Sea snakes include >60 species in two major clades, *Hydrophis* and *Aipysurus-Emydocephalus*, which shared a semi-aquatic common ancestor ∼6-18 Mya and exhibit highly contrasting evolutionary histories since their transitions from terrestrial to marine species (Sanders et al. 2013; Lee et al. 2016; Nitschke et al. 2018). Both of these lineages have independently developed adaptations to the aquatic environment including valve-like nostrils allowing for full closure when underwater and tail paddles for efficient underwater movement (Lillywhite 2014). However, the *Aipysurus-Emydocephalus* lineage has continued to evolve at the same rate as terrestrial lineages of Hydrophiinae, diverging into 9 species, while the Hydrophis lineage has rapidly radiated into 48 species (Sanders et al. 2010). Following major changes in habitat ecology, such as sea snakes’ transition from a terrestrial to a marine habitat, organisms must adapt to their new environment, with transposons potentially contributing to adaptations (Schlötterer et al. 2015; Marques et al. 2018). Here we report an analysis of transposons in sea snake genomes, where the marine environment appears to have fostered the repeated, independent acquisition of these transposons through horizontal transfer of transposons (HTT). The repeated HTT suggests that direct effects of the environment on genome structure may be an important but overlooked driver of evolutionary change during major ecological transitions.

## Results

### Annotation of sea snake transposons

We previously performed ab initio repeat annotation of the olive sea snake (*Aipysurus laevis*) genome (Ludington et al., in prep) using CARP (Zeng et al. 2018) and RepeatModeler (Smit and Hubley) to compare its repetitive content to that of its terrestrial relatives *Notechis scutatus* (tiger snake) and *Pseudonaja textilis* (eastern brown snake). Most repetitive sequences identified by CARP were not well classified because both CARP and RepeatModeler rely on homology to reference sequences from Repbase (Bao et al. 2015), a database of repeats from highly studied species that are evolutionarily distant to Hydrophiinae. This reliance on sequence homology for *ab initio* repeat annotation of newly sequenced species often results in the incorrect annotation of repeats (Platt et al. 2016).

We used a structural homology approach based on the presence a variety of protein domains in these poorly annotated repeats to identify subfamilies of LINEs, Penelople and LTR retrotransposons, endogenous retroviruses, and Tc1/Mariner DNA transposons. All the TE subfamilies identified in the *A. laevis* genome were also present in *P. textilis* and *N. scutatus* with the exception of five LINE subfamilies (see Figure 1). Two of the five LINEs subfamilies, Rex1-Snek_1 (five full-length copies found) and Rex1-Snek_2 (three full-length copies found) belong to the CR1/Jockey superfamily but share less than 100 bp nucleotide sequence homology. Manual curation of a multiple sequence alignment of the five full-length copies identified by CARP revealed Rex1-Snek_1 to be three subfamilies; henceforth named Rex1-Snek_1, Rex1-Snek_3 and Rex1-Snek_4. Rex1-Snek_3 and Rex1-Snek_4 have 90% and 89% pairwise identity with Rex1-Snek_1 respectively. The other three subfamilies, RTE-Snek_1 (three full-length sequences found), RTE-Snek_2 (one full-length sequence found) and Proto2-Snek (one full-length sequence found) belong to the RTE superfamily but have no significant nucleotide sequence homology. In addition to the full-length sequences, we identified hundreds of highly similar copies with 5’ truncation patterns characteristic of recently active LINEs (Figure 2). Specifically, coverage plots of the RTE-Snek_1, RTE-Snek_2 and Proto2-Snek families are typical of LINEs, with a clear pattern of 5’ truncated insertions (Luan et al. 1993). All seven LINE subfamilies were most similar to Repbase TE reference sequences from a marine annelid worm, a marine crustacean and teleost fishes (Bao et al. 2015) (see Table 1, SI Dataset 1).

**Table 1.** Most similar RepBase and curated repeats for each repeat family in species outside of snakes. RepBase was searched using the seven consensus *Aipysurus laevis* repeats using relaxed BLASTN parameters (see Methods). A database of our curated repeats from all searched species (see Methods) was searched using the seven consensus *Aipysurus laevis* repeats using default BLASTN parameters.

| Most similar RepBase sequences |  |  |  |
| --- | --- | --- | --- |
| Repeat (query) | Species (target repeat) | Percent identity | Hit length (bp) |
| Rex-Snek_1 | *Petromyzon marinus* (Rex1-1_PM) | 67.5 | 1,359 |
| Rex-Snek_2 | *Cyprinus carpio* (Rex1-1_CCa) | 75.9 | 2,795 |
| Rex-Snek_3 | *Petromyzon marinus* (Rex1-1_PM) | 66.7 | 1,359 |
| Rex-Snek_4 | *Petromyzon marinus* (Rex1-1_PM) | 64.2 | 2,796 |
| RTE-Snek_1 | *Petromyzon marinus* (RTE-2_PM) | 62.9 | 3,100 |
| RTE-Snek_2 | *Chrysemys picta* (RTE-9_CPB) | 65.3 | 2,926 |
| Proto2-Snek | *Oryzias latipes* (Proto2-1_OL) | 65.6 | 666 |
| Most similar curated repeats |  |  |  |
| Rex-Snek_1 | *Oryzias latipes* | 85.0 | 2,987 |
| Rex-Snek_2 | *Miichthys miiuy* | 78.7 | 2,594 |
| Rex-Snek_3 | *Oryzias latipes* | 82.2 | 2,973 |
| Rex-Snek_4 | *Oryzias latipes* | 81.6 | 2,960 |
| RTE-Snek_1 | *Laticauda colubrina* | 84.9 | 3,252 |
| RTE-Snek_2 | *Hippocampus comes* | 74.4 | 3184 |
| Proto2-Snek | *Epinephelus lanceolatus* | 75.4 | 3,299 |

**Figure 1.**
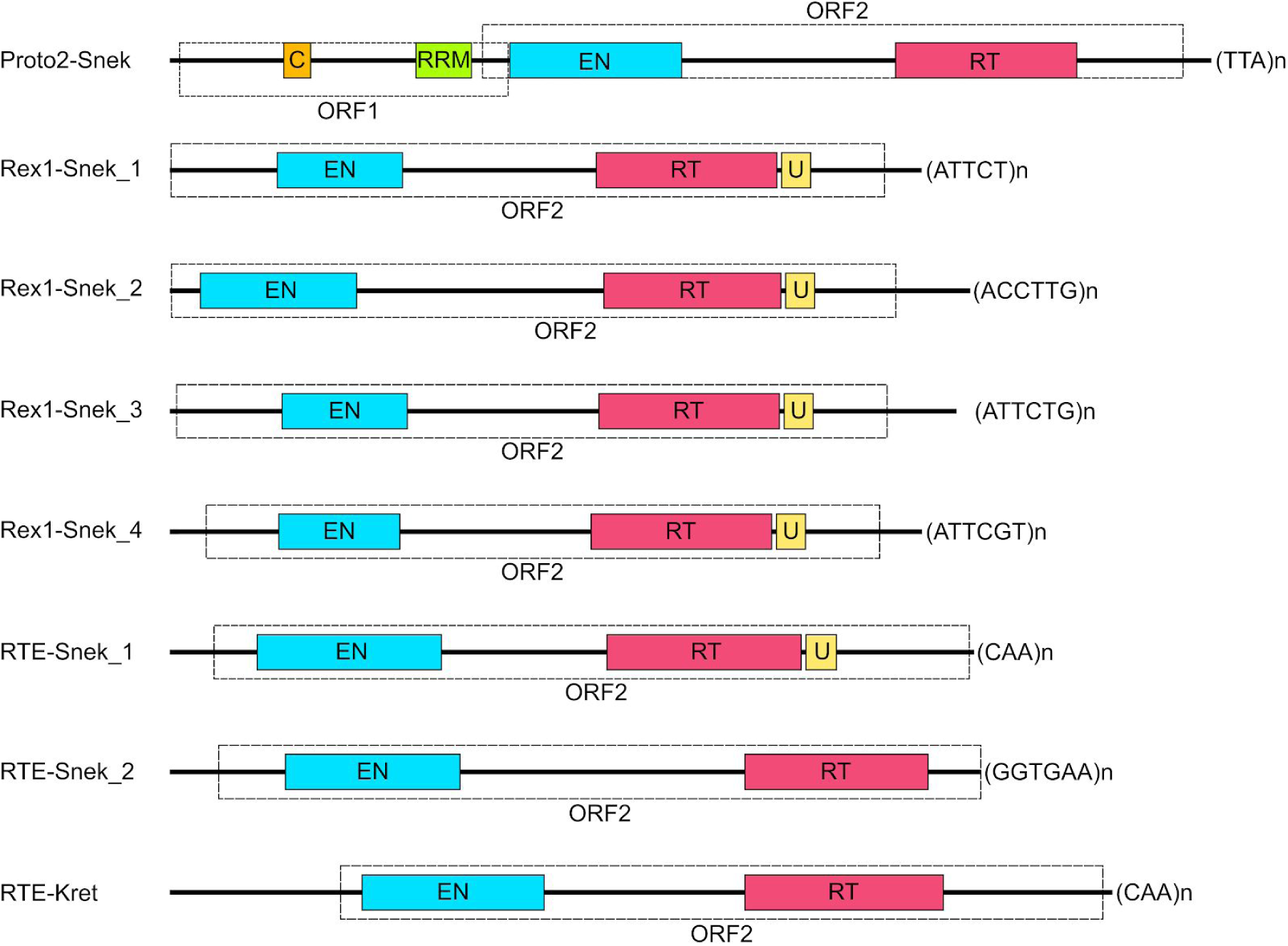
Structure of the seven horizontally transferred LINEs in *Aipysurus laevis* - Proto2-Snek, Rex1-Snek_1, Rex1-Snek_2, Rex1-Snek_3, Rex1-Snek_4, RTE-Snek_1 and RTE-Snek_2; and the one horizontally transferred LINE in *Laticauda colubrina* - RTE-Kret. Cyan represents endonuclease (EN), red reverse transcriptase (RT), orange coiled coil (CC), green RNA-recognition motif (RRM), and yellow domain of unknown function 1891 (U). Protein domains were identified using RPSBLAST (Marchler-Bauer et al. 2017) and HHpred (Zimmermann et al. 2018) searches against CDD and Pfam (Finn et al. 2016; Marchler-Bauer et al. 2017) databases and the coiled coil domain was identified using PCOILS (Gruber et al. 2006).

**Figure 2.**
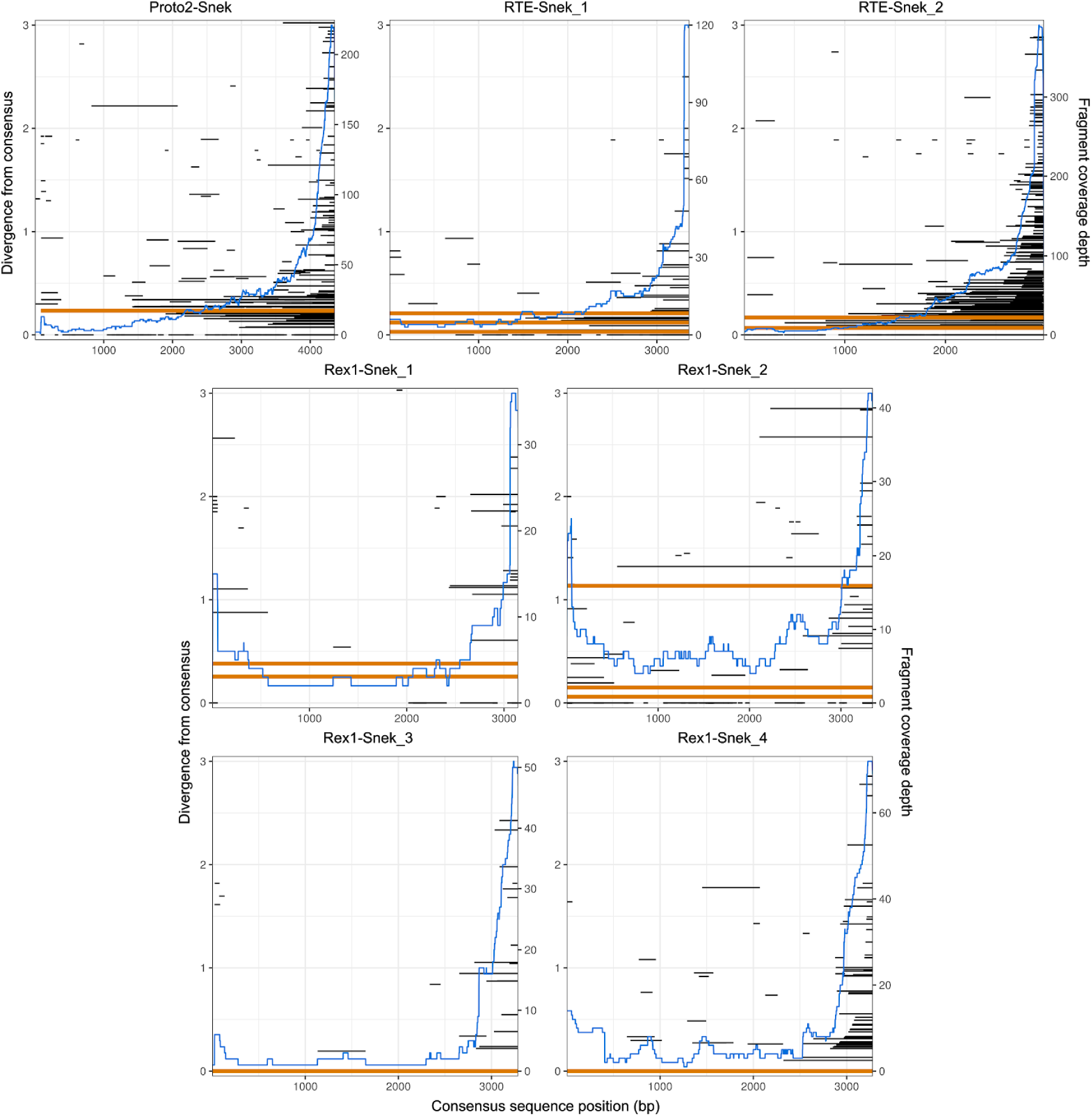
Coverage and divergence from consensus of the seven horizontally transferred LINEs identified in the *Aipysurus laevis* genome. LINE fragments were identified with BLASTN (Altschul et al. 1990; Camacho et al. 2009) and plotted using ggplot2 (Wickham 2011) using the consensus2genome script (https://github.com/clemgoub/consensus2genome). The blue line represents the depth of coverage of fragments aligned to the subfamily consensus sequence (shown on right hand Y-axis). Each horizontal line represents the divergence of a fragment and its position mapped to the repeat consensus (position shown on X-axis); red shows full length repeats and black shows repeat fragments. The divergence from consensus of the repeats is shown on left hand Y-axis.

The absence of these recently active LINE subfamilies from terrestrial snakes that diverged within the last approximately 18 Mya, combined with the finding that they were most similar to LINEs from distantly related aquatic organisms, suggested HTT as a likely explanation. There are three diagnostic features of HTT: 1) the sporadic presence of a TE family within a set of closely related species, 2) a higher than expected degree of sequence identity in long diverged species and 3) discordant topologies for the phylogenies of transposons and their host species (Silva et al. 2004).

### Presence/absence in closely related species

As mentioned above, the seven LINEs are absent from close terrestrial relatives of *A. laevis*. To test if the LINEs have a sporadic distribution in nearer relatives we performed reciprocal BLAST+ nucleotide searches for their presence in two closely related sea snake genome assemblies, *Hydrophis melanocephalus* (slender-necked sea snake) and *Emydocephalus ijimae* (Ijima’s turtleheaded sea snake); the two closely related terrestrial species, *N. scutatus* and *P. textilis*; an independently aquatic species, *Laticauda colubrina* (yellow-lipped sea krait); and a distant terrestrial relative, *Ophiophagus hannah* (king cobra). The reciprocal search for RTE-Snek_1 revealed a similar yet distinct RTE subfamily present in *L. colubrina*, henceforth referred to as RTE-Kret. From these searches, we found RTE-Snek_1 was restricted to *A. laevis* and RTE-Kret to be restricted to *L. colubrina*. In addition to being present in *A. laevis*, Proto2-Snek was also present in *E. ijimae*; Rex1-Snek_1, Rex1-Snek_2 and RTE-Snek_2 in *E. ijimae* and *H. melanocephalus*; and Rex1-Snek_3 and Rex1-Snek_4 in *H. melanocephalus*. This reciprocal search confirmed all seven LINEs to be absent from both terrestrial (*N. scutatus, P. textilis* and *O. hannah*) and aquatic (*L. colubrina*) outgroups, and RTE-Kret to be restricted to *L. colubrina* (Fig. 3, SI Fig. 3-10).

**Figure 3.**
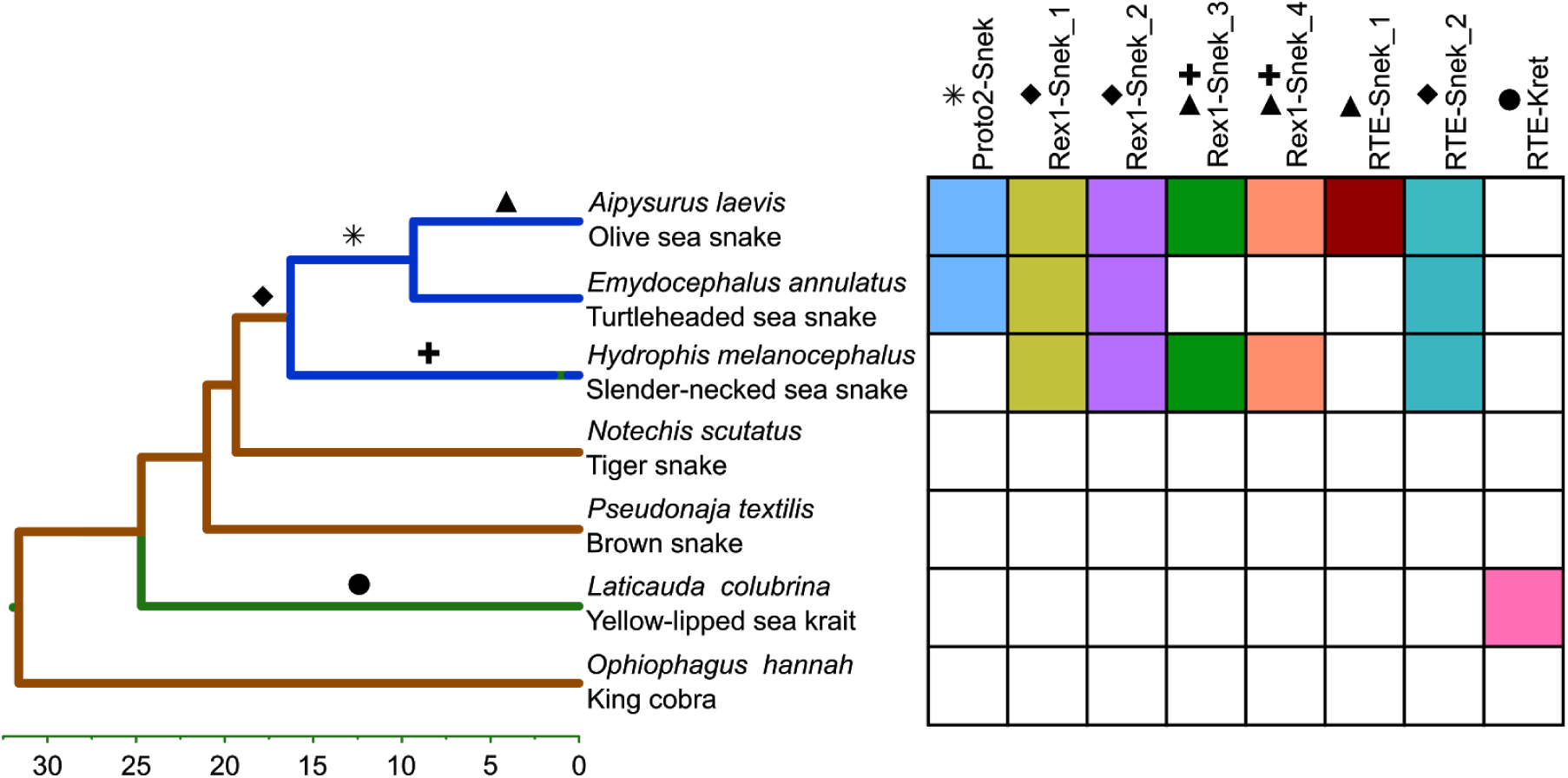
Presence of the eight HTT LINE subfamilies across the phylogeny of elapid snakes (adapted from Lee et al, 2016 (Lee et al. 2016)). Colour of lineage represents habitat - marine species are blue, terrestrial brown and amphibious green. Each symbol represents the likely timing of horizontal transfers. Presence/absence determined using reciprocal BLASTN search (Altschul et al. 1990; Camacho et al. 2009) using default parameters.

As an independent verification of presence/absence and to look for potential current activity of the LINEs, we searched transcriptomes of a variety of tissues from three sea snakes - *A. laevis, A. tenuis* and *Hydrophis major* (SI Dataset 2). We identified high-identity fragments of all four Rex1-Sneks and RTE-Snek_2 in at least one tissue of *A. laevis, A. tenuis* and *H. major*. High identity transcript fragments of RTE-Snek_1 and Proto2-Snek were present in *A. laevis* and *A. tenuis*, yet largely absent from all *H. major* tissues, with only one small RTE-Snek_1-like fragment present in a *H. major* testis transcriptome. The presence of LINE transcripts both confirmed the presence of specific LINEs and indicated potential ongoing retrotransposition of these elements. The presence/absence of all seven LINEs identified in *A. laevis* across other sea snakes and their close terrestrial relatives is indicative of multiple, independent HTT events (Fig. 3).

### Search for HTT donor species

In order to identify potential donor taxa for the seven LINEs transferred into sea snakes, we searched for and curated similar LINEs in more than 600 metazoan genomes (SI Dataset 3). Our manual curation found homologous Rex1s in fish and squamates, Proto2s in fish, and RTEs widespread across a variety of marine organisms including fish, echinoderms, corals and sea kraits (see Fig. 4, SI Dataset 4). We then aligned our original LINE sequences against a database containing both our curated repeats and Repbase repeats. All seven of our original LINEs were most similar to curated LINEs found in marine species (Table 1) with pairwise identity for all closest hits between 75-85%. Rex1-Snek_1, Rex1-Snek_2, Rex1-Snek_3 and Rex1-Snek_4 were most similar to Rex1s curated from a variety of fish genomes. Proto2-Snek was most similar to a Proto2 from the European carp (*Cyprinus carpio*) genome and RTE-Snek_1 most similar to RTE-Kret from *L. colubrina*. However, we were unable to identify plausible donor species as none of the cross species alignments were greater than 87% nucleotide sequence identity (Table 1).

**Figure 4.**
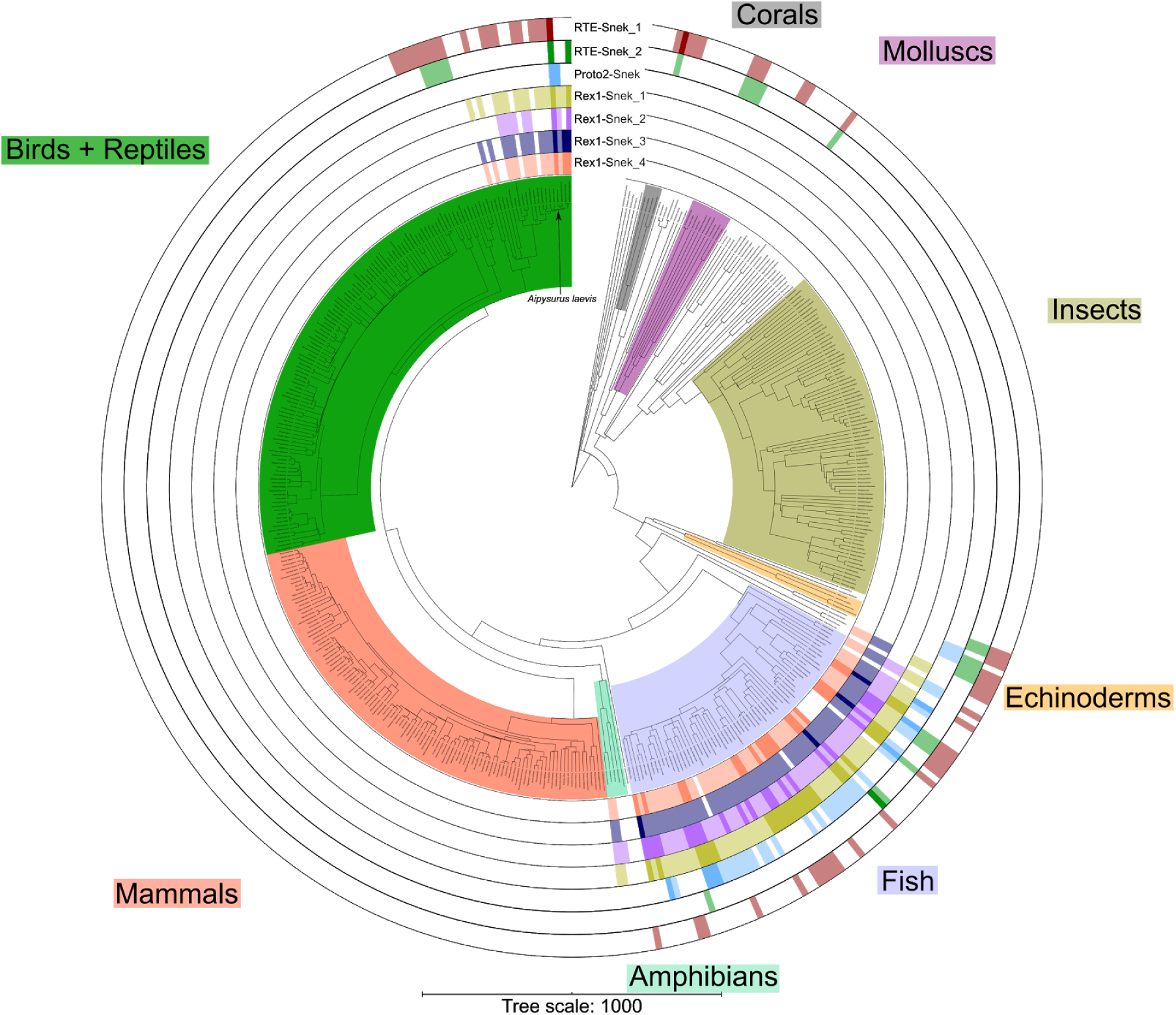
Presence of the seven HTT LINEs across 458 Metazoa. In each ring darker shading represents the presence of at least one sequence over 1000 bp in length showing 75% or higher pairwise identity to the LINE, lighter shading represents the presence of at least one sequence over 1000 bp with less than 75% pairwise identity, and white represents the complete absence of similar sequences. Presence of LINEs identified using BLASTN (Altschul et al. 1990; Camacho et al. 2009) and plotted in iToL (Letunic and Bork 2019). Species tree generated using TimeTree (Hedges et al. 2006), manually edited to correct elapid phylogeny to fit (Lee et al. 2016). Interactive tree available at https://itol.embl.de/shared/jamesdgalbraith.

### Discordant phylogenies of RTEs and of Rex1s compared to host species

As extreme discordance between repeat and species phylogenies is evidence of HTT, we compared the tree topology of all RTEs, Proto2s and Rex1s, using both Repbase sequences and our curated sequences, to the species tree topology. As illustrated in figure 5, the species and repeat phylogenies of all seven sea snake LINEs and the *Laticauda* LINE are highly discordant, evidenced by their clustering with teleost fishes, confirming likely HTT events from marine organisms into sea snakes and sea kraits. The presence/absence pattern observed, illustrated in figure 3, suggests 6 to 8 transfers into sea snakes and 1 into sea kraits.

**Figure 5.**
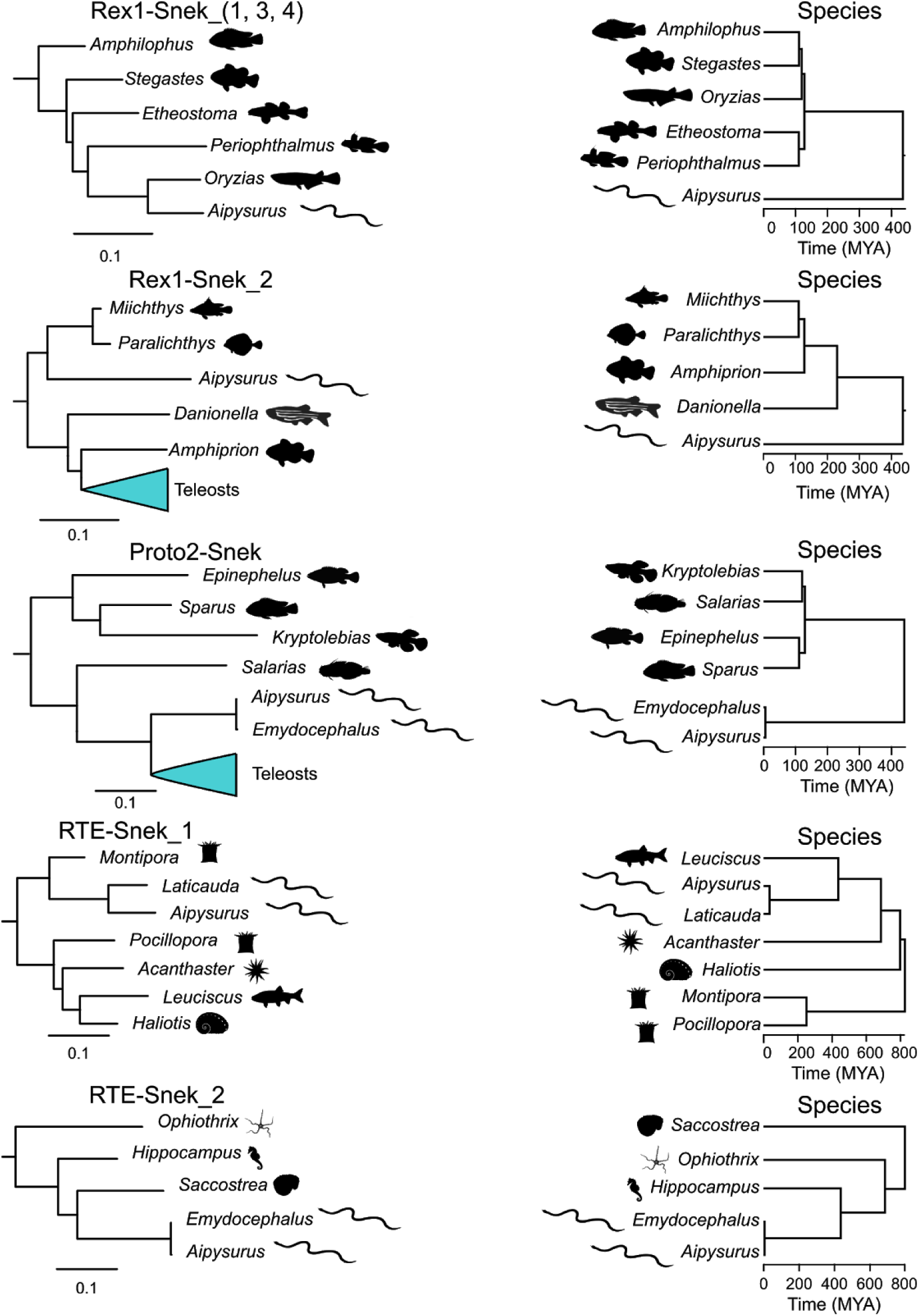
Excerpts from the phylogenies of all intact curated and RepBase RTE-like LINEs and all intact curated and RepBase Rex1s compared to host species phylogeny. The blue triangles on the left represent condensed subtrees of LINE sequences. TE phylogeny scale bar represents substitutions per site. Extracts from larger phylogenies constructed using RAxML (Stamatakis 2014) based on MAFFT (Katoh and Standley 2013) alignments trimmed with Gblocks (Talavera and Castresana 2007) (for full phylogenies see SI Appendix, Fig. S1 and Fig. S2). Species trees constructed with TimeTree (Hedges et al. 2006).

### Insertions in and near coding regions

To identify any insertions of these LINEs in *A. laevis* which might have altered gene expression or protein structure, we identified all insertions in or near regions annotated as genes, exons and 5’ UTRs (SI Table 1). To test for potential assembly errors that might have yielded erroneous insertions near genes, we searched for orthologous insertions and flanking regions in two other sea snake assemblies from closely related species; *E. ijimae* and *H. melanocephalus*. Intersects of our genome and repeat annotations from Ludington et al. (unpublished) initially revealed 19 insertions of HTT LINEs within 5000 bp of 5’ UTRs, one into a coding exon and 3 into 3’ UTRs.

However, 8 insertions may be the result of assembly errors in *A. laevis*, as their flanking sequences were in the middle of two different contigs in both *E. ijimae* and *H. melanocephalus*. The remaining insertions were confirmed as present solely in *A. laevis*. We report these 15 insertions in Table 2, but testing of these insertions for adaptive significance will have to await improvement of the genome assembly and population genetic data for *A. laevis*. We note that many of these genes are likely to have pleiotropic effects as regulators of transcription or protein turnover, thus complicating future assessments of their adaptive significance.

**Table 2:** HTT LINEs inserted into exons or within 5000 bp upstream of 5’ UTRs of genes within the *A. laevis* assembly.

| Gene | LINE | Distance to 5' UTR (bp) | Insertion size (bp) |
| --- | --- | --- | --- |
| Acetyl-CoA Acyltransferase 1 (*ARIH1*) | Proto2-Snek | 223 | 85 |
| KN Motif And Ankyrin Repeat Domains 4 (*KANK4*) | Proto2-Snek | 4987 | 161 |
| Potassium Calcium-Activated Channel Subfamily N Member 4 (*KCNN4*) | Proto2-Snek | 3746 | 98 |
| Outer Mitochondrial Membrane Lipid Metabolism Regulator OPA3 (*OPA3*) | Rex1-Snek_2 | 3149 | 81 |
| Rabaptin, RAB GTPase Binding Effector Protein 1 (*RABEP1*) | Proto2-Snek | 1389 | 99 |
| Valosin Containing Protein Lysine Methyltransferase (*VCPKMT*) | Rex1-Snek_1 | 512 | 76 |
| Cdc42 effector protein 4 (*CDC42EP4*) | RTE-Snek_2 | 1475 | 422 |
| Gamma-aminobutyric acid receptor subunit alpha-3 (*GABRA3*) | RTE-Snek_2 | 4247 | 95 |
| Leucine-zipper-like transcriptional regulator 1 (*LZTR1*) | RTE-Snek_2 | 2066 | 421 |
| Polyadenylate-binding protein 2 (*PABPN1*) | RTE-Snek_2 | 145 | 431 |
| Parvalbumin alpha (*PVALB*) | RTE-Snek_2 | 4152 | 52 |
| Deaminated glutathione amidase (*NIT1*) | RTE-Snek_2 | In coding exon | 228 |
| Adenylate cyclase type 4 (*ADCY4*) | RTE-Snek_2 | In 3' UTR | 130 |

## Discussion

We provide strong evidence that the seven LINEs identified in *A. laevis* were horizontally transferred to sea snakes after their transition to a marine habitat, from marine species. This is based on the absence of all seven LINEs from terrestrial relatives and the discordance of the species and LINE phylogenies (Figs. 3 and 5). While all seven are currently expressed based on transcriptome data, the number of large, near-identical fragments of RTE-Snek_1, RTE-Snek_2 and Proto2-Snek found within the *A. laevis* genome is larger than for Rex1s and indicates potentially greater replication since the original insertion events that occurred at most 12 Mya.

As all seven of the HTT LINEs are most similar to LINEs in distantly related marine metazoans, the donor species for each is likely a fish or marine invertebrate. However, the degree of sequence divergence between the LINE from *L. colubrina* and the seven LINEs from *A. laevis* from the most similar LINEs from aquatic species means we cannot identify a donor species. As very few species with ranges overlapping that of *Laticauda* and *Aipysurus* have been sequenced and the range of *Aipysurus* spans highly biodiverse habitats, it is unlikely we will further narrow the donor of any of these seven LINEs without significant additional genome sequence data from Indo-Pacific tropical marine species.

While we were unable to determine the donor species, our finding of HTT between marine species is in line with multiple past studies finding HTT within and across marine phyla. HTT is prolific and particularly well described in aquatic microbial communities (reviewed in-depth in Sobecky and Hazen, 2009 (Sobecky and Hazen 2009)). HTT of LINEs, LTR retrotransposons and DNA transposons has been reported in marine metazoans, with past studies describing the transfer of Rex1s and Rex3s between teleost fishes (Volff et al. 2000; Carducci et al. 2018), *Steamer*-like LTR retrotransposons both within and across phyla (Metzger et al. 2018), L1 and BovB LINEs within and across phyla (Ivancevic et al. 2018), and Mariner DNA transposons between diverse crustaceans (Casse et al. 2006). What sets our findings apart is that HTTs have occurred multiple times as a result of the recent terrestrial to marine transition of the *Aipysurus*/*Hydrophis* common ancestor. The transfer of all seven LINEs occurred <18 Mya from aquatical animal donor species that diverged from snakes >400 Mya (Broughton et al. 2013; Hughes et al. 2018). As illustrated in figure 3, the varying presence/absence of the seven LINEs across the three species of sea snakes is indicative of multiple HTT events as opposed to a single event. The recent timing of HTT into marine squamates is not specific to sea snakes, as we found transfer of an RTE-Kret to the sea kraits which underwent an independent transition to the same habitat. These repeated invasions suggest the marine environment potentially fosters HTT, with more examples likely to be revealed by additional genome sequences from marine species.

The likely ongoing replication of all 6 *A. laevis* HTT LINEs, as evidenced by both the presence of insertions and transcripts with near 100% identity, continues to contribute genetic material to the evolution of *Aipysurus*. Previous investigators have reported entire genes, exons, regulatory sequences and noncoding RNAs in vertebrates derived from transposons, as well as TE insertions leading to genomic rearrangement (reviewed in-depth in Warren et al. (Warren et al. 2015)). For example, the insertion of CR1 fragments near phospholipase A2 venom genes in vipers led to non-allelic homologous recombination, in turn causing duplication of these genes (34). Rapid genomic innovation would have been necessary for *Aipysurus* to adapt to the marine environment, with the independent evolution of paddle-like tails, salt excretion glands and dermal photoreception following their divergence from their most recent common ancestor with *Hydrophis* (Brischoux et al. 2012; Sanders et al. 2012; Crowe-Riddell et al. 2019). Other adaptations are likely to have occurred or are occurring for sea snakes to better adapt to their marine habitat, as evolutionary transitions from terrestrial to marine habits entail massive phenotypic changes spanning metabolic, sensory, locomotor, and communication-related traits. Whilst we have identified some HTT LINE insertions near genes that may potentially alter gene expression, future research to examine the association between these HTT-derived sequences and adaptation will have to await improvement of the *A. laevis* genome assembly and population genomic data.

## Conclusions

Our findings reveal repeated HTT of LINEs into a fully marine lineage following their transition from a terrestrial environment, while Australian terrestrial elapids show no evidence of HTT of LINEs. In addition, we discovered one homologous HTT event to have occurred in a lineage of snakes which has independently transitioned to a semi-aquatic environment, providing more evidence that the marine environment promotes HTT. The continued expression of all seven LINEs also suggests ongoing impact on the evolution of *Aipysurus*. Combined, this supports a likely role for habitat transitions making a direct contribution to the evolution of metazoan genome content, rather than genomes evolving solely in response to selection imposed by changing environmental conditions. We can view the ancestral genome and novel selection from a new habitat as the two “parents” that give rise to new species, but our data indicate that HTT from the new environment may act as a “third parent”, with this more likely in some habitats.

## Materials and Methods

### Identification and classification of repetitive sequences in *Aipysurus laevis*

We identified repetitive sequences present in the Ludington et al. (unpublished) *A. laevis* assembly using CARP (Zeng et al. 2018). Using RPSTBLASTN 2.7.1+ (Marchler-Bauer and Bryant 2004) and a custom library of position specific scoring matrices from the CDD and Pfam databases (Finn et al. 2016; Marchler-Bauer et al. 2017) we identified protein domains present in all repetitive sequences over 800 bp in length found by CARP. We treated consensus sequences containing over 80% of both an exo-endonuclease domain and a reverse transcriptase domain as potential LINEs. We used CENSOR 4.2.29 (Kohany et al. 2006) to classify the consensus sequences. To reduce redundancy, we aligned all potential LINEs to all other potential LINEs using BLASTN 2.7.1+ (Altschul et al. 1990; Camacho et al. 2009) with default parameters and constructed clusters based on pairwise identity (97% or higher). From each cluster the longest sequence was treated as the representative sequence.

To create a better consensus for each LINE subfamily, we manually curated new consensus sequences using a “search, extend, align, trim” method. Using the largest consensus from a subfamily we used BLASTN 2.7.1+ (Altschul et al. 1990; Camacho et al. 2009) with default parameters to search for the repeat within the *A. laevis* genome. We selected the best thirty hits over 1000 bp based on bitscore and extended the coordinates of these sequences by 1000 bp at each end of the hit. We constructed multiple sequence alignments (MSAs) of the extended sequences using MAFFT v7.310 (Katoh and Standley 2013). Where multiple full length sequences showing significant non-homology were present the LINE subfamily was split into multiple subfamilies. Finally, we manually edited the extended sequences in Geneious Prime 2020.0.2 to remove non-homologous regions and created a new consensus sequence. If only one full length copy of a subfamily was present in the genome it was used as the consensus sequence. We used PCOILs (Gruber et al. 2006) and HHpred (Zimmermann et al. 2018) searches of the translated ORFs against the CDD and Pfam databases (Finn et al. 2016; Marchler-Bauer et al. 2017) to identify any additional protein domains or structures present in the seven LINEs.

### Search for LINEs in closely related species genomes and transcriptomes

To determine if the seven LINEs were present in closely related species we used BLASTN 2.7.1+ (Altschul et al. 1990; Camacho et al. 2009) to perform a reciprocal nucleotide search of appropriate elapid snake genomes downloaded from GenBank (Benson et al. 2017). After discovering and curating RTE-Kret in *L. colubrina* the process was repeated. Similarly, we used BLASTN 2.7.1+ (Altschul et al. 1990; Camacho et al. 2009) to perform reciprocal searches for the seven LINEs in transcriptomes from various tissues of *A. laevis, A. tenuis* and *H. major* from Crowe-Riddell et al. (Crowe-Riddell et al. 2019).

### Search for and curation of similar LINEs in other metazoan genomes

To identify other species containing the seven LINEs, we used BLASTN 2.7.1+ (Altschul et al. 1990; Camacho et al. 2009) to search over 600 metazoan genomes downloaded from Genbank (44) using relaxed parameters (-evalue 0.00002 -reward 3 -penalty -4 -xdrop_ungap 80 -xdrop_gap 130 -xdrop_gap_final 150 -word_size 10 -dust yes -gapopen 30 -gapextend 6). We treated species containing a hit of at least 1000 bp as potentially containing a similar LINE. In species potentially containing a similar LINE we attempted to manually curate the LINE using a variant of the “search, extend, align, trim” method described above. We used a consensus of the initial hits within the species as the query for the BLASTN search of the genome, and extended hits by 3000 bp in each direction. If an MSA appeared to contain multiple LINE families, the MSA was split into the families and consensuses constructed for each individual family. To identify LINEs in *A. laevis* similar to those identified in other elapid snakes we used the same approach as above, using LINEs curated from *N. scutatus* as the query. The same method was used to curate RTE-Kret from the *L. colubrina* assembly.

### Characterising divergence patterns in the HT repeats across Hydrophiinae

To identify fragments of the seven HTT LINEs identified in *A. laevis* and determine their divergence from the consensus sequences we performed a reciprocal best hit search using BLASTN 2.7.1+ (Altschul et al. 1990; Camacho et al. 2009) on the *A. laevis, E. ijimae, H. cyanocinctus, H. melanocephalus, N. scutatus, P. textilis, L. colubrina* and *O. hannah* assemblies. Following the identification and curation of RTE-Kret this was repeated.

### Repeat phylogeny construction

For constructing repeat phylogenies we created two libraries; one containing all curated and Repbase Rex1s and another containing all curated and Repbase RTE-like (Proto2, RTE and BovB) LINEs. In addition, each library contained an outgroup LINE based on the Eickbush and Malik (Eickbush and Malik 2002) phylogeny of LINEs. We removed all sequences not containing at least 80% of both the endonuclease and reverse transcriptase domains from each library based on RPSTBLASTN+ (Marchler-Bauer and Bryant 2004) searches against the NCBI CDD (Marchler-Bauer et al. 2017).

We created MSAs of each library of LINEs using MAFFT v7.310 (Katoh and Standley 2013) and removed poorly aligned regions using Gblocks (Talavera and Castresana 2007). Finally we constructed phylogenies from the trimmed MSA using RAxML (Stamatakis 2014) with 20 maximum likelihood trees and 500 bootstraps.

### Species phylogeny construction

We used TimeTree (Hedges et al. 2006) to infer species phylogenies presented in Figure 4. In cases in which a species of interest was not present in the TimeTree database, where possible we used an appropriate species from the same clade in its place and corrected the species names on the resulting tree.

### Repeat insertions near and in genes

Using the plyranges (Lee et al. 2019) and GenomicRanges R packages (Lawrence et al. 2013) (53, 54), we identified any insertions of the HTT LINEs into coding exons, UTRs and upstream of 5’ UTRs as annotated in the repeat annotation and gene annotations from Ludington et. at. (unpublished) (see overlapSearch.R script in github repository).

To confirm insertions were assembled correctly we used BLASTN+ to search for the repeats extended by 2000 bp in each direction in the *E. ijimae* and *H. melanocephalus* assemblies. We selected the best hits from each species based on query coverage and percent identity. Using MAFFT v7.310 (Katoh and Standley 2013) we constructed multiple sequence alignments of each extended repeat and the corresponding regions from the two other assemblies (see insertionConfirmation.R in github repository). By manually viewing the resulting alignment as well and the raw BLASTN+ output we determined if the repeat insertions were assembled correctly.

All scripts used are available at https://github.com/jamesdgalbraith/HT_Workflow

## Acknowledgments

We thank Dan Kortschak (University of Adelaide), Jesper Boman (Uppsala University), Valentina Poena (Uppsala University), Atma Ivancevic (University of Colorado) and Ed Chuong (University of Colorado) for helpful suggestions and discussions.

