## Supplementary Information for "New environment, new invaders - repeated horizontal transfer of LINEs to sea snakes"

* David L. Adelson[1]

**This PDF file includes:**

SI Figures 1-10
SI Table 1
Legends for Datasets S1 to S8

**Other supplementary materials for this manuscript include the following:**

Datasets S1 to S8

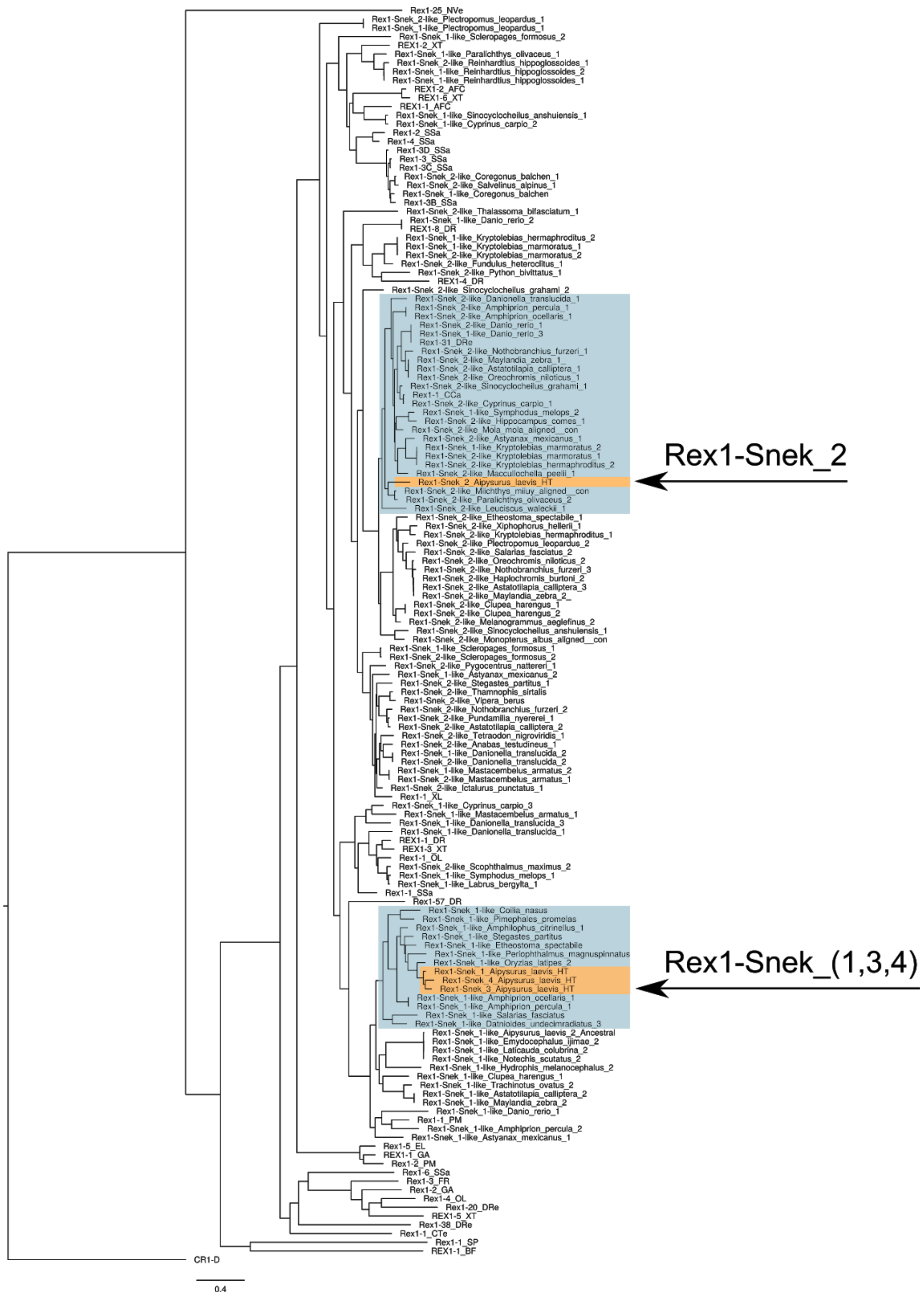


**Figure S1.** Full tree of all curated and Repbase Rex1-like LINEs containing both an endonuclease and a reverse transcriptase domain. Horizontally transferred LINEs in sea snakes are phylogenetically discordant as seen by their  highlighting in brown, within the most similar LINEs found in teleosts  in blue. Phylogeny created using RAxML (Stamatakis 2014) from a multiple sequence alignment generated using MAFFT (Katoh and Standley 2013) and trimmed using Gblocks (Talavera and Castresana 2007). Sequences available in SI Dataset 6, Newick tree in SI Dataset 7.

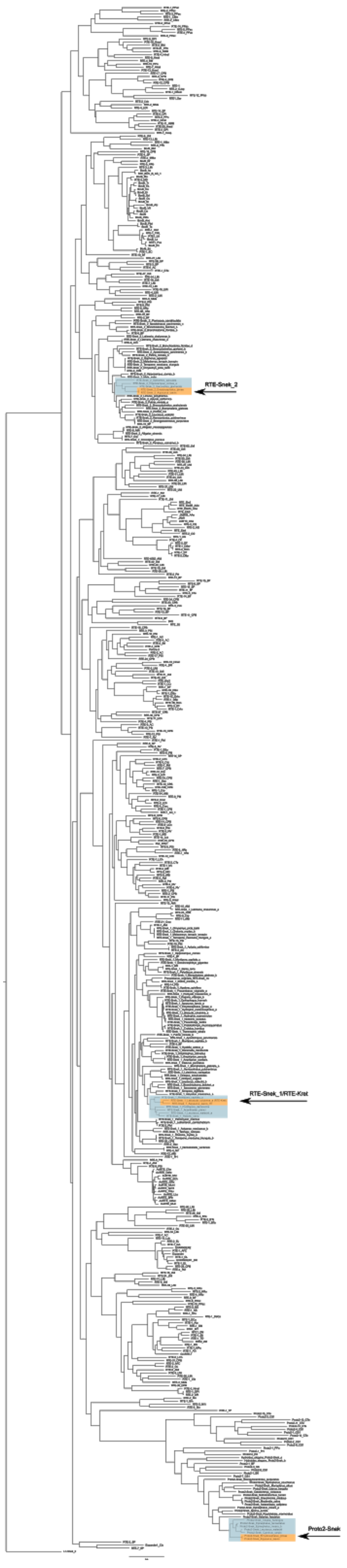
RTE-Snek_2
RTE-Snek_1/RTE-Kret
Proto2-Snek

**Figure S2.** Full tree of all curated and Repbase RTEs and Proto2s containing both an endonuclease and a reverse transcriptase. Horizontally transferred LINEs in sea snakes are phylogenetically discordant as seen by their highlighting in brown, within the most similar LINEs found in lamprey, corals, bivalve, starfish, fishes  in blue. Phylogeny created using RAxML (Stamatakis 2014) from a multiple sequence alignment generated using MAFFT (Katoh and Standley 2013) and trimmed using Gblocks (Talavera and Castresana 2007). Sequences available in SI Dataset 6, Newick tree in SI Dataset 8.

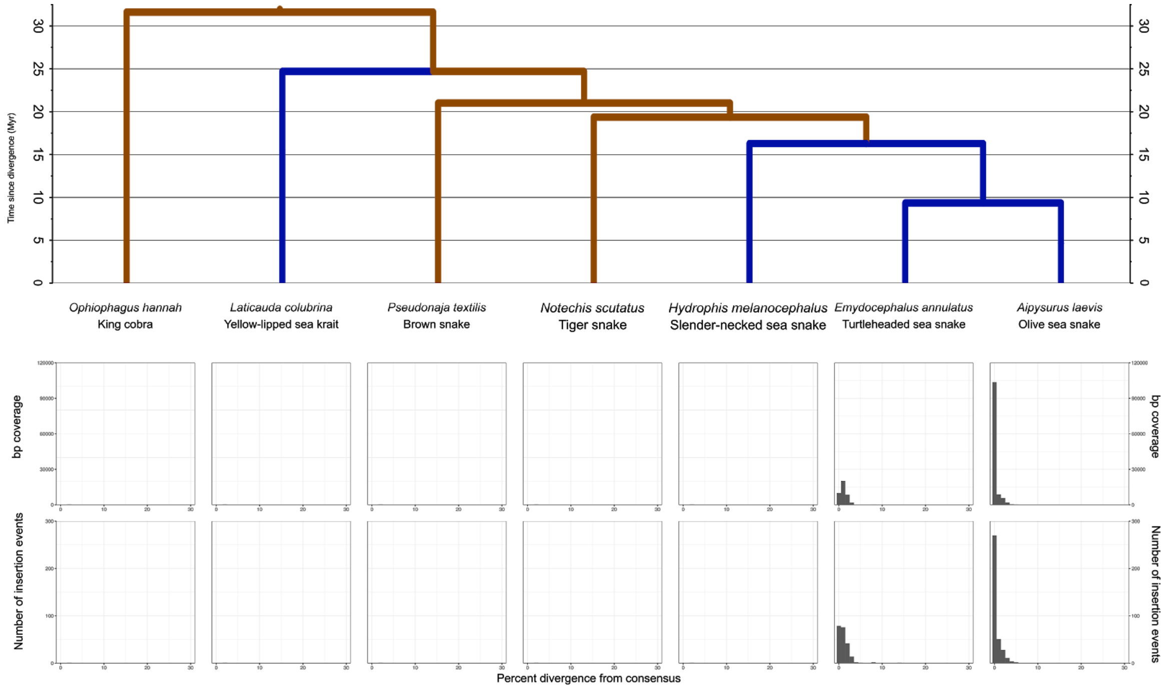


**Figure. S3.** Number of insertions using fragments as a proxy, and bp coverage vs divergence from consensus of Proto2-Snek insertions identified in *Ophiophagus hannah*, *Laticauda colubrina* and six Hydrophiinae. Presence of LINEs detected using BLASTN+ 2.7.1 (Altschul et al. 1990; Camacho et al. 2009) and plotted in RStudio (RStudio Team 2015) using ggplot2 (Wickham 2011).

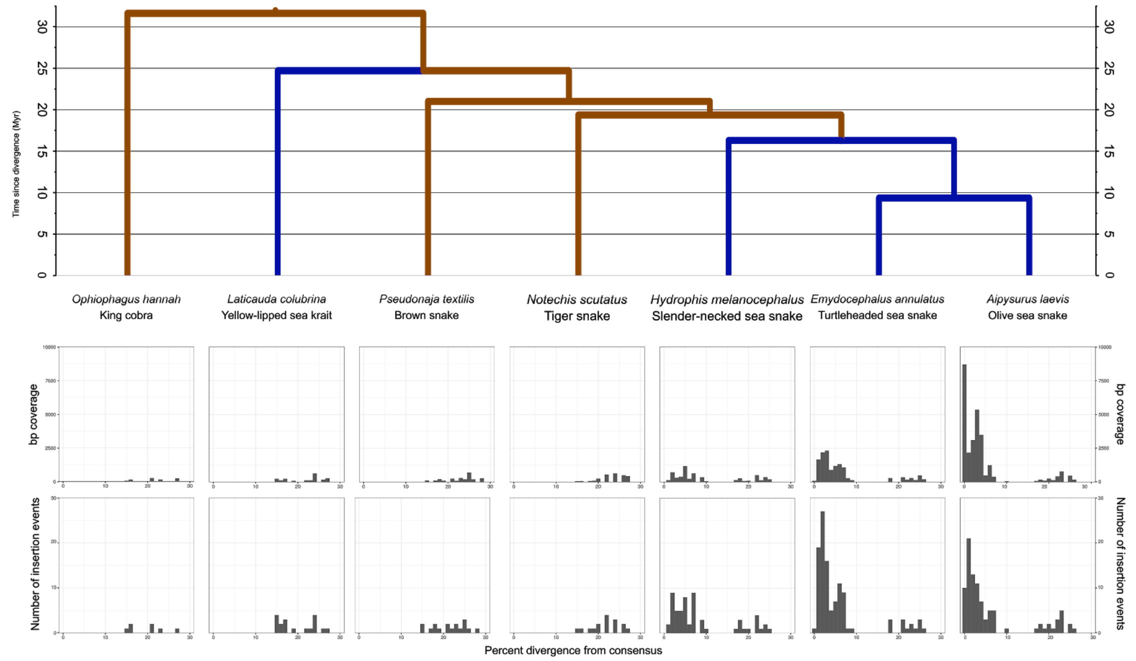


**Figure S4.** Number of insertions using fragments as a proxy, and bp coverage vs divergence from consensus of Rex1-Snek_1 insertions identified in *Ophiophagus hannah*, *Laticauda colubrina* and six Hydrophiinae. Note that insertions with >15% divergence represent similarity to distant, ancestral Rex1 elements found in non-avian reptiles. Presence of LINEs detected using BLASTN+ 2.7.1 (Altschul et al. 1990; Camacho et al. 2009) and plotted in RStudio (RStudio Team 2015) using ggplot2 (Wickham 2011).

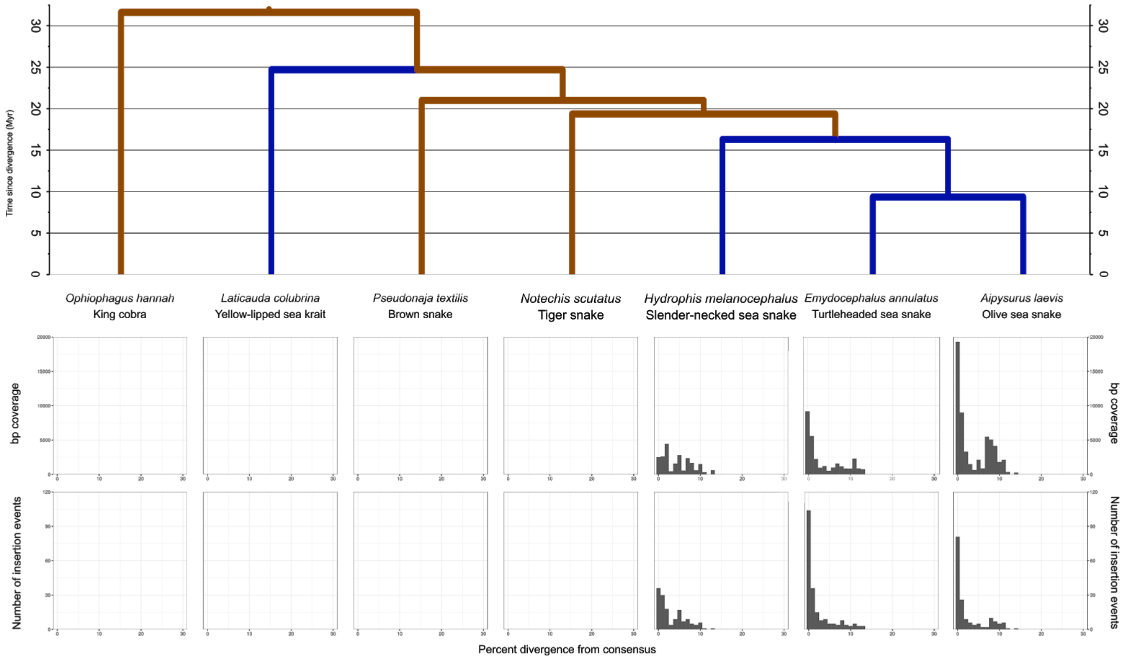


**Figure S5.** Number of insertions using fragments as a proxy, and bp coverage vs divergence from consensus of Rex1-Snek_2 insertions identified in *Ophiophagus hannah*, *Laticauda colubrina* and six Hydrophiinae. Presence of LINEs detected using BLASTN+ 2.7.1 (Altschul et al. 1990; Camacho et al. 2009) and plotted in RStudio (RStudio Team 2015) using ggplot2 (Wickham 2011).

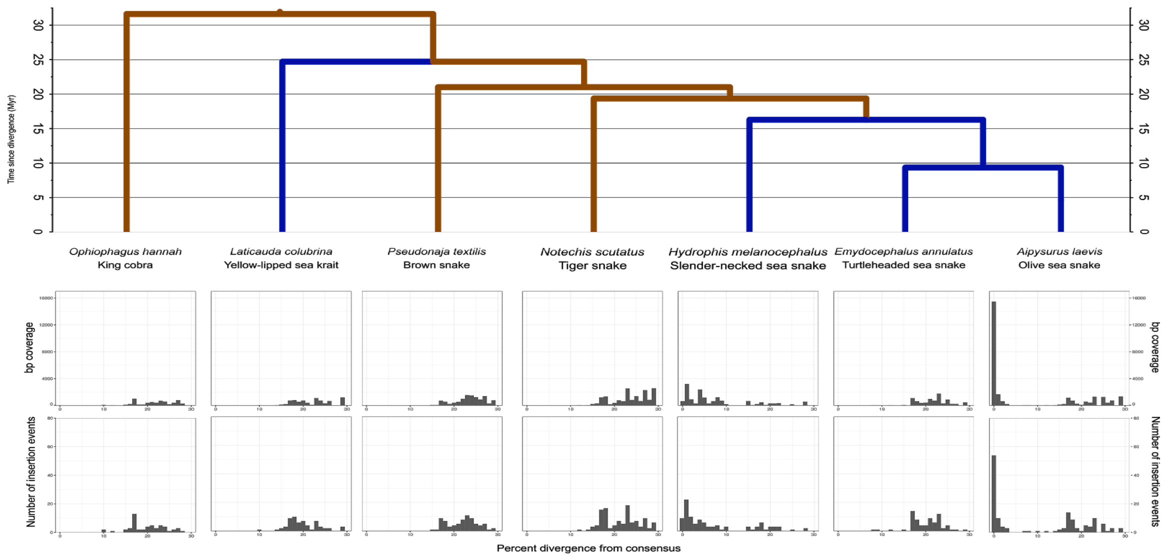


**Figure S6.** Number of insertions using fragments as a proxy, and bp coverage vs divergence from consensus of Rex1-Snek_3 insertions identified in *Ophiophagus hannah*, *Laticauda colubrina* and six Hydrophiinae. Note that insertions with >15% divergence represent similarity to distant, ancestral Rex1 elements found in non-avian reptiles. Presence of LINEs detected using BLASTN+ 2.7.1 (Altschul et al. 1990; Camacho et al. 2009) and plotted in RStudio (RStudio Team 2015) using ggplot2 (Wickham 2011).

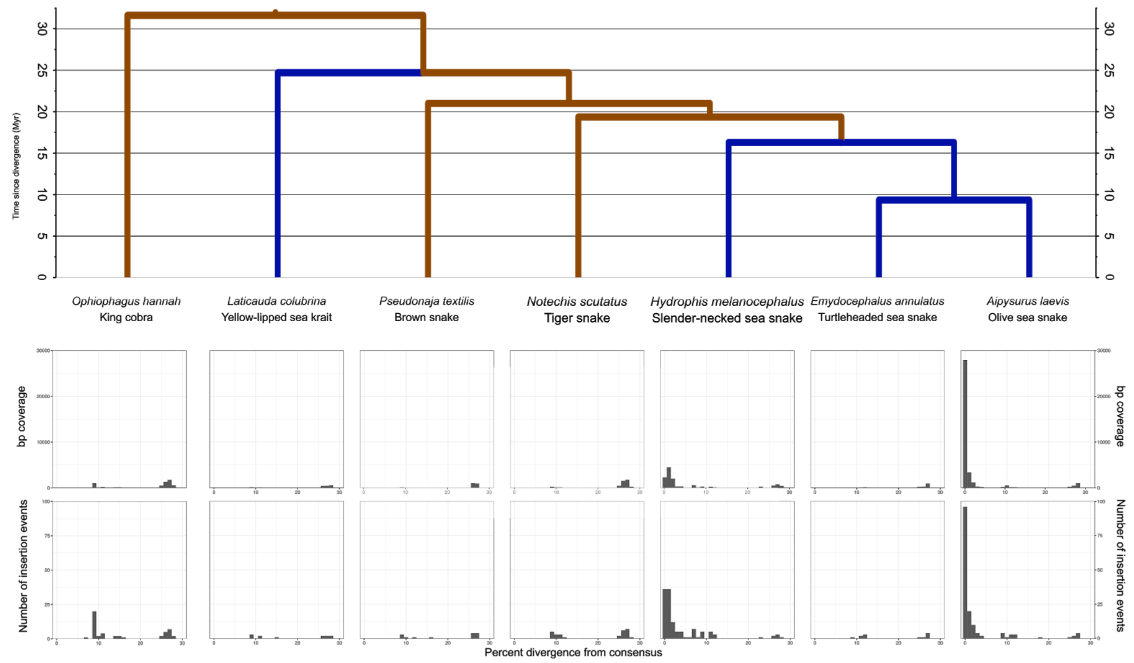


**Figure S7.** Number of insertions using fragments as a proxy, and bp coverage vs divergence from consensus of Rex1-Snek_4 insertions identified in *Ophiophagus hannah*, *Laticauda colubrina* and six Hydrophiinae. Note that insertions with >15% divergence represent similarity to distant, ancestral Rex1 elements found in non-avian reptiles. Presence of LINEs detected using BLASTN+ 2.7.1 (Altschul et al. 1990; Camacho et al. 2009) and plotted in RStudio (RStudio Team 2015) using ggplot2 (Wickham 2011).

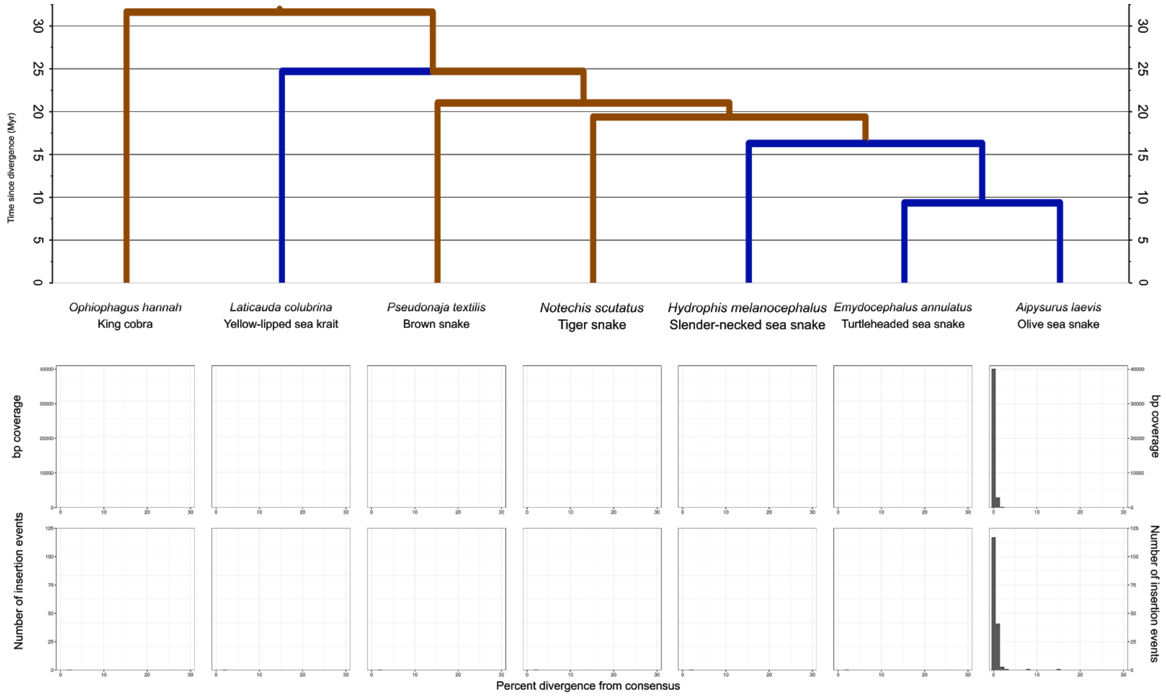


**Figure S8.** Number of insertions using fragments as a proxy, and bp coverage vs divergence from consensus of RTE-Snek_1 insertions identified in *Ophiophagus hannah*, *Laticauda colubrina* and six Hydrophiinae. Presence of LINEs detected using BLASTN+ 2.7.1 (Altschul et al. 1990; Camacho et al. 2009) and plotted in RStudio (RStudio Team 2015) using ggplot2 (Wickham 2011).

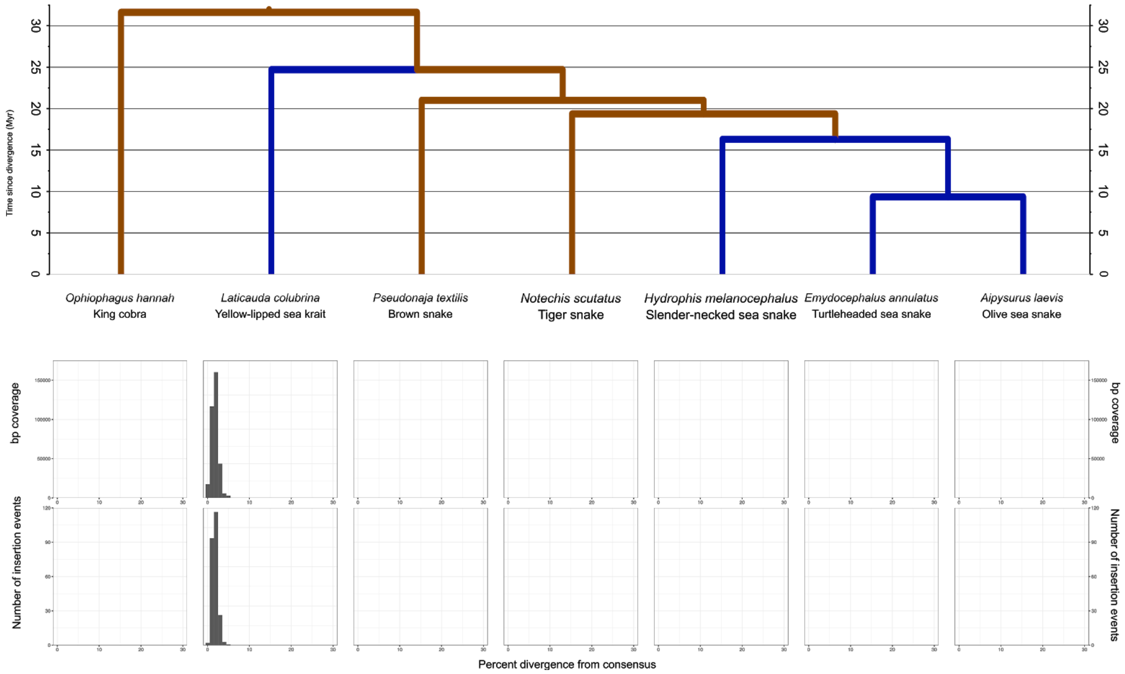


**Figure S9.** Number of insertions using fragments as a proxy, and bp coverage vs divergence from consensus of RTE-Kret insertions identified in *Ophiophagus hannah*, *Laticauda colubrina* and six Hydrophiinae. Presence of LINEs detected using BLASTN+ 2.7.1 (Altschul et al. 1990; Camacho et al. 2009) and plotted in RStudio (RStudio Team 2015) using ggplot2 (Wickham 2011).

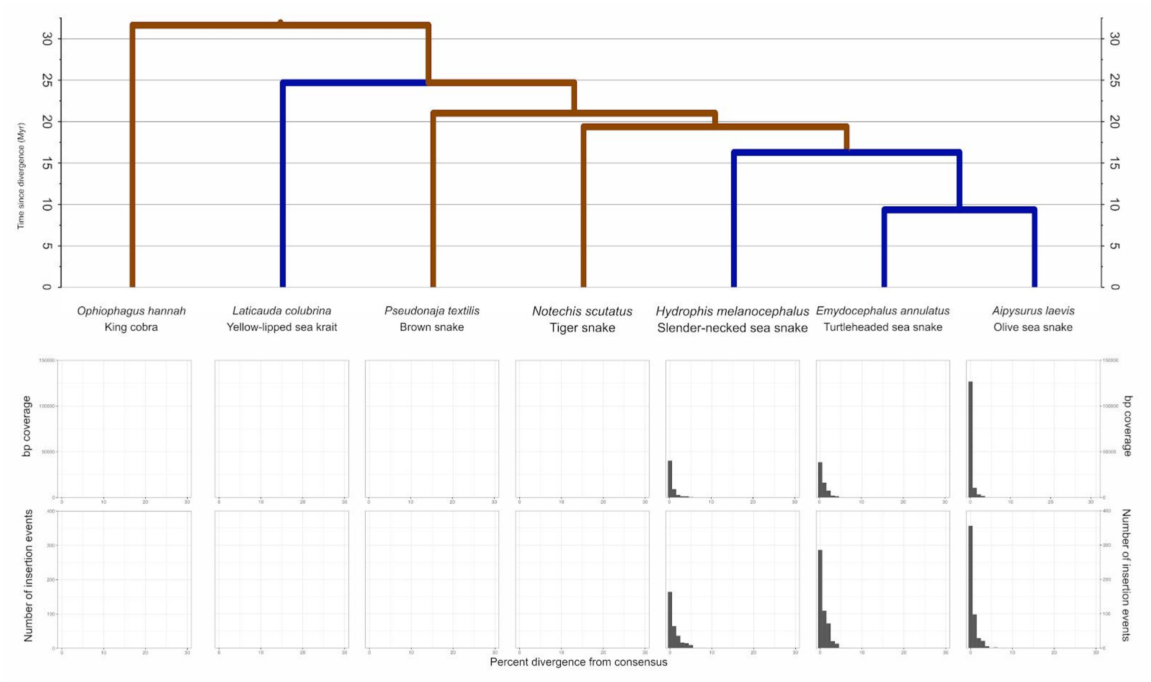


**Figure S10.** Number of insertions using fragments as a proxy, and bp coverage vs divergence from consensus of RTE-Snek_2 insertions identified in *Ophiophagus hannah*, *Laticauda colubrina* and six Hydrophiinae. Presence of LINEs detected using BLASTN+ 2.7.1 (Altschul et al. 1990; Camacho et al. 2009) and plotted in RStudio (RStudio Team 2015) using ggplot2 (Wickham 2011).

| | HTT LINEs | All LINEs |
|---|---|---|
| Total insertions | 1478 | 139942 |
| Insertions on same scaffold as genes | 536 | 43033 |
| Total Insertions into genes | 76 | 8491 |
| Insertions into coding exons | 1 | 725 |
| Insertions into UTRs | 2 | 572 |
| Insertions within 5000bp of 5’ UTR | 22 | 4022 |

**Table S1.** Raw number of all LINE insertions and HT LINE insertions into *A. laevis* genome, exons and upstream of 5’UTRs with a divergence from consensus below 10%. Insertions detected using Intersects of genome and repeat annotations of *A. laevis* assembly from Ludington et al. unpublished in RStudio (RStudio Team 2015) using GenomicRanges and plyranges (Lawrence et al. 2013; Lee et al. 2019).

**Dataset S1.** Searches for all seven HT LINEs in RepBase (Bao et al. 2015) using CENSOR 4.2.29 (Kohany et al. 2006).

**Dataset S2.** BLASTN+ 2.7.1 (Altschul et al. 1990; Camacho et al. 2009) searches for all six repeats in 11 sea snake transcriptomes from Crowe-Riddell. Results below 250bp in length and <95% identity discarded.

**Dataset S3.** Latin species names and versions of all public genomes used. All were downloaded from GenBank (48).

**Dataset S4.** Number of repeats similar to Rex1-Snek_1, Rex1-Snek_2, RTE-Snek and Proto2-Snek found in all species search using BLASTN+ 2.7.1 (Altschul et al. 1990; Camacho et al. 2009) with relaxed parameters (see Methods).

**Dataset S5.** Multi-FASTA of the six horizontally transferred LINEs found in *Aipysurus laevis* and the one horizontally transferred LINE found in *Laticauda colubrina.*

**Dataset S6.** Multi-FASTA of all manually curated Rex1s, RTEs and Proto2s.

**Dataset S7.** Newick tree of the four Rex1s found in *Aipysurus laevis*, all manually curated Rex1s and all Rex1s from RepBase. Phylogeny created using RAxML (Stamatakis 2014) from a multiple sequence alignment generated using MAFFT (Katoh and Standley 2013) and trimmed using Gblocks (Talavera and Castresana 2007).

**Dataset S8.** Newick tree of Proto2-Snek, RTE-Snek, RTE-Kret, all manually curated Rex1s and all Rex1s from RepBase. Phylogeny created using RAxML (Stamatakis 2014) from a multiple sequence alignment generated using MAFFT (Katoh and Standley 2013) and trimmed using Gblocks (Talavera and Castresana 2007)
